# Pentose phosphate pathway metabolites are induced and required for nucleotides by latent Kaposi’s Sarcoma infection of endothelial cells

**DOI:** 10.64898/2026.11.23.768784

**Authors:** Jacqueline Alexander, Michael Lagunoff

## Abstract

Kaposi’s Sarcoma Herpesvirus (KSHV) is the etiologic agent of Kaposi’s Sarcoma (KS). As a herpesvirus, KSHV has both latent and lytic phases and is predominantly found in the latent phase in KS spindle cells, the main proliferating cell of KS tumors. KS spindle cells express markers of endothelium. We previously found that latent KSHV infection of endothelial cells dramatically alters host cell metabolism including increased glucose uptake and glycolysis. The Pentose Phosphate Pathway (PPP), a shunt from glycolysis is often increased in cancer cells and some lytic viral infections. Here we found that KSHV increases intermediates of the oxidative half of the PPP in tert-immortalized microvascular endothelial cells. Chemical inhibition or knockout of an enzyme in the oxidative half of the PPP demonstrated the essential nature of the PPP during latent infection of endothelial cells. However, inhibition of the non-oxidative half has no effect on latently infected cells. The oxidative half of the PPP produces ribose-5-phosphate a substrate for nucleotide synthesis. We found that both purine and pyrimidine synthesis are necessary for survival of KSHV latently infected endothelial cells but not for mock infected cells. Addition of exogenous ribose recovers much of the cell death induced by inhibition of the PPP, indicating that the PPP appears to be required for nucleotide synthesis. The requirement of nucleotide synthesis in latently infected cells raises the possibility that the PPP and nucleotides are needed for more than rapid viral replication during viral infection.

**Importance:** Treatment of KS tumors relies upon cancer chemotherapy which targets cancer cells but not latent KSHV infection potentially leading to high relapse rates in KS patients treated with chemotherapy. Identifying KSHV host dependencies during latent infection may provide novel treatment approaches for latent infection. We demonstrate that the PPP and nucleotide synthesis are required for the survival of endothelial cells latently infected with KSHV providing new approaches to target latently infected cells.

## Introduction

Kaposi sarcoma (KS), an angioproliferative tumor, is still among the most frequent AIDS-associated malignancies worldwide and is one of the most common cancers in central and southeast Africa where it affects both individuals living with and without HIV/AIDS (1–3). KS is a highly vascularized tumor which predominately consists of spindle cells that express markers of endothelium (4–10). Kaposi sarcoma herpesvirus (KSHV), also known as human herpesvirus-8 (HHV-8) is an oncogenic ɣ-herpesvirus (1) that is the etiologic agent of Kaposi sarcoma (KS) and two B-cell lymphoproliferative diseases, primary effusion lymphoma (PEL) and multicentric Castleman’s diseases (MCD) (11, 12). As a herpesvirus, KSHV has both a lytic and latent state. Lytic viral infection occurs with rapid viral replication and cell death while in the latent viral state the virus is maintained as an episome without death of the cell. KS spindle cells are primarily latently infected with KSHV, though a low proportion of cells express markers of lytic replication (13–15). Infection of both cultured primary and TERT-immortalized microvascular endothelial cells can recapitulate infection rates that are seen in KS spindle cells (16, 17). During KSHV latency only a limited number of viral genes and microRNAs are expressed making it difficult to target latent infection (15, 18, 19). While treatment regimens are dependent on the severity and spread of the disease and the patient’s immune system, the most prevalent treatment for KS is chemotherapy (20). Relapses are common in KS patients treated with chemotherapy as these approaches target cancer cells but do not directly target latently infected cells (21, 22). Newer approaches to target latency are necessary. However, since latent herpesviruses are difficult to directly target, targeting latent KSHV host cell dependencies may provide novel treatment approaches.

Previous work from our lab has examined how KSHV infection alters host metabolism. We identified significant changes to the host cell during lytic replication of endothelial cells and surprisingly, during latent KSHV infection of endothelial cells as well (23, 24). From a metabolomics screen of KSHV latently infected endothelial cells and uninfected cells, we identified several metabolic pathways that were altered during KSHV latency in endothelial cells including three major metabolic pathways that are also common in cancer: glycolysis, glutaminolysis and fatty acid synthesis (24–26). We have shown that latent KSHV upregulates glucose uptake and glycolysis which leads to an increase in lactate production (25). All of this occurs with a reduction in the use of oxidative phosphorylation showing that KSHV induces the Warburg effect, a phenomenon that is seen in most, if not all, cancer cells where glycolysis becomes the preferred energy source, even in the presence of oxygen (27, 28). Latent infection also induces glutamine uptake and upregulates a glutamine transporter, SLC1A5, through the transcription factor c-Myc and its dimerization proteins (26). Additionally, KSHV infection increased more than half of the measured long-chain fatty acids in our metabolomics study.

Importantly, when KSHV latently infected endothelial cells are treated with inhibitors to different steps of any of these three major metabolic pathways, it leads to cell death specifically in latently infected cells (24–26). KSHV infected primary effusion lymphomas, a KSHV dependent B-cell cancer, also exhibited differences in similar metabolic pathways (29).

The pentose-phosphate pathway (PPP) is a metabolic pathway that branches off from glycolysis. This pathway produces both nicotinamide adenine dinucleotide phosphate (NADPH), and ribose 5-phosphate (R5P), a precursor sugar for the synthesis of nucleotides (30, 31). This pathway has been shown to be important in cancer cells as NADPH is necessary as a reducing agent for reactive oxygen species (ROS) to maintain the low ROS levels necessary to stimulate cell proliferation (32). Cancer cells also need ribose-5-phosphate to increase nucleotide synthesis to enable the proliferation of new cancer cells (33).

Here we show that intermediates of the pentose phosphate pathway (PPP) are increased in KSHV latently infected endothelial cells. Inhibition or deletion of the rate limiting enzyme in the oxidative PPP, G6PD, leads to cell death specifically in latently infected cells showing that the PPP is necessary for the survival of latently infected cells. Inhibition of the non-oxidative PPP enzyme, transketolase, does not similarly induce cell death in latently infected cells indicating the latter part of the non-oxidative half of the PPP, where metabolites can reenter glycolysis, is not necessary. Inhibition of purine and pyrimidine nucleotides leads to cell death during KSHV latency showing that nucleotides are necessary for the survival of latently infected endothelial cells. Addition of exogenous ribose leads to rescue of the cell death that we see with inhibiting the PPP. Therefore, latent KSHV infection induces the PPP to produce the precursor metabolite, ribose-5-phosphate, for nucleotide synthesis pathways during KSHV latency.

## Results

### Latent KSHV Infection of Endothelial Cells Induces the Pentose Phosphate Pathway

We previously performed a global metabolomics screen to identify changes to host cell metabolism induced by KSHV. The metabolomic screen identified that nearly a third of 194 detectable metabolites were altered by 96 hours post KSHV infection in Tert-Immortalized Microvascular Endothelial (TIME) cells (24). Analysis of intermediates of the Pentose phosphate pathway (PPP) indicate that three out of the four detectable PPP metabolites in our metabolomics: 6-phosphogluconate, ribose-5-phosphate, and ribulouse-5-phoshate were at elevated levels in KSHV-infected cells in comparison to mock infected cells (Figure 1A and B). Intracellular levels of 6-phosphogluconate were elevated by 1.4-fold in KSHV infected cells at 48 hours post infection and increased to 3.2-fold by 96 hours post infection. Ribose 5-phosphate was elevated to 1.6-fold at 48 hours post infection while, ribulose 5-phosphate was elevated to 1.8-fold, with the caveat that this metabolite cannot be distinguished from xylulose 5-phosphate in this analysis (Figure 1B). These results indicate that metabolites in the PPP are increased in KSHV latently infected TIME cells (Figure 1B).

**Figure 1:**
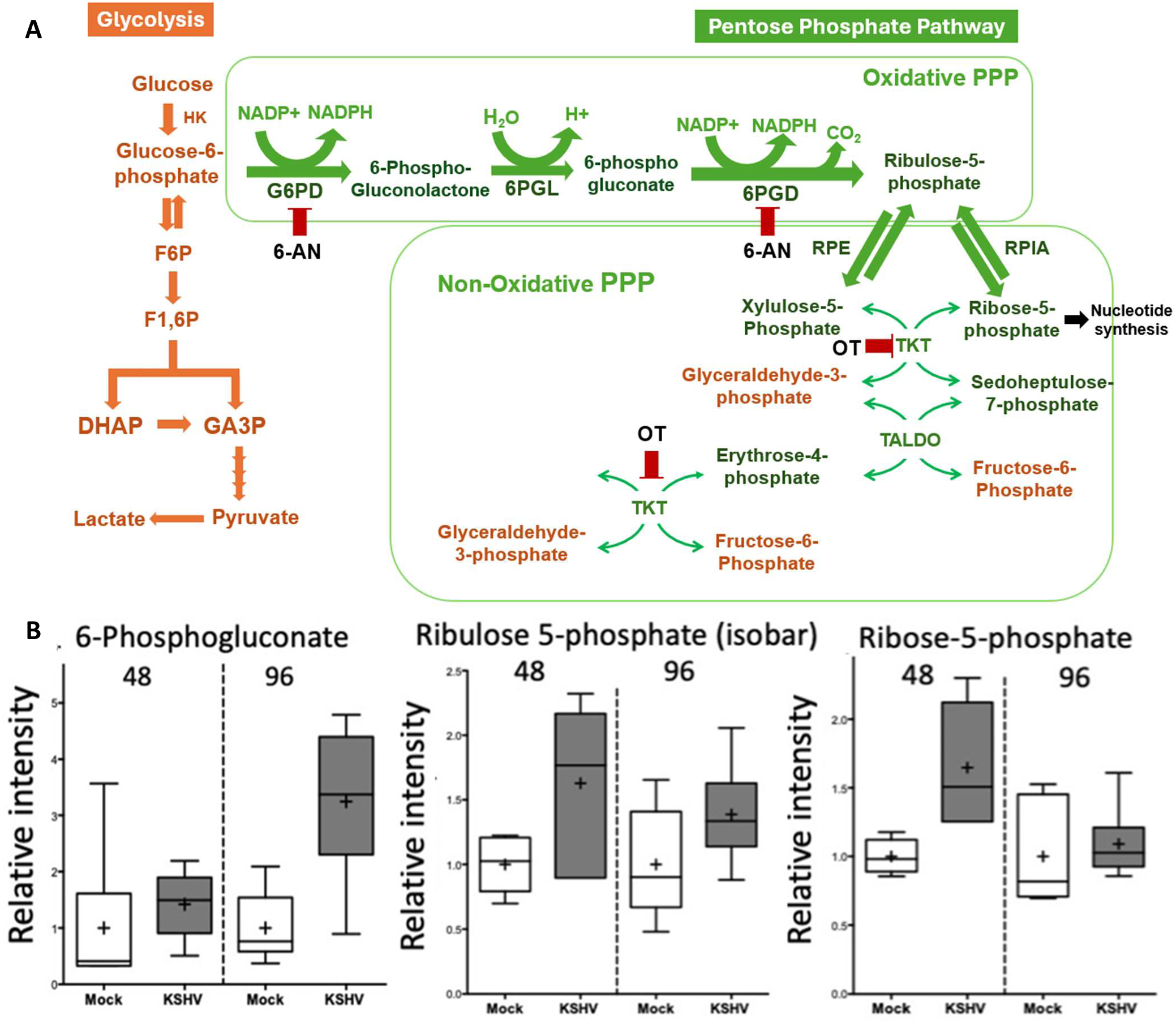
Pentose Phosphate pathway metabolites, 6-phosphogluconate, ribulose 5-phosphate and ribose 5-phosphate, are at elevated levels in latently infected KSHV endothelial cells. (A) Schematic of glycolysis and the pentose phosphate pathway (PPP). Orange represents glycolysis while green represents the PPP. Red markers represent drug inhibitors for specific enzymes in the PPP. (B) Box and whisper plot of average intensity of PPP metabolites determined by global metabolomics screen of Mock and KSHV infected TIME cells at 48 and 96 hours post infection. Mock infected cells are represented by white boxes while KSHV infected cells are represented in grey. Measurements were taken from 6 biological replicates. The average of each group is represented by a (+) sign. P values ≤ 0.05.

### The PPP is required for survival of KSHV latently infected endothelial cells

To determine if the PPP plays a role during KSHV latency, we examined the effect of 6-Aminonicotinamide (6-AN), a competitive inhibitor of glucose-6-phosphate dehydrogenase (G6PD), the first and rate-limiting enzyme of the PPP (40, 41) (Figure 1A). TIME cells were either mock or KSHV infected and at 48 hours post infection, when cells have established latency, both mock and KSHV infected cells were treated with 6-AN or the vehicle control for an additional 48 hours. Prior to inhibitor treatment, cells were collected at 48 hours post infection for an immunofluorescence assay (IFA) to confirm that greater than 88% of infected TIME cells were expressing LANA, a KSHV latent protein. Post treatment with 6-AN, cells were harvested from each condition and accessed for cell death rates via a trypan blue exclusion assay. KSHV-infected TIME cells were significantly more sensitive to 6-AN showing an average 21% percent cell death in comparison to Mock cells’ 6% cell death (Figure 2A).

**Figure 2:**
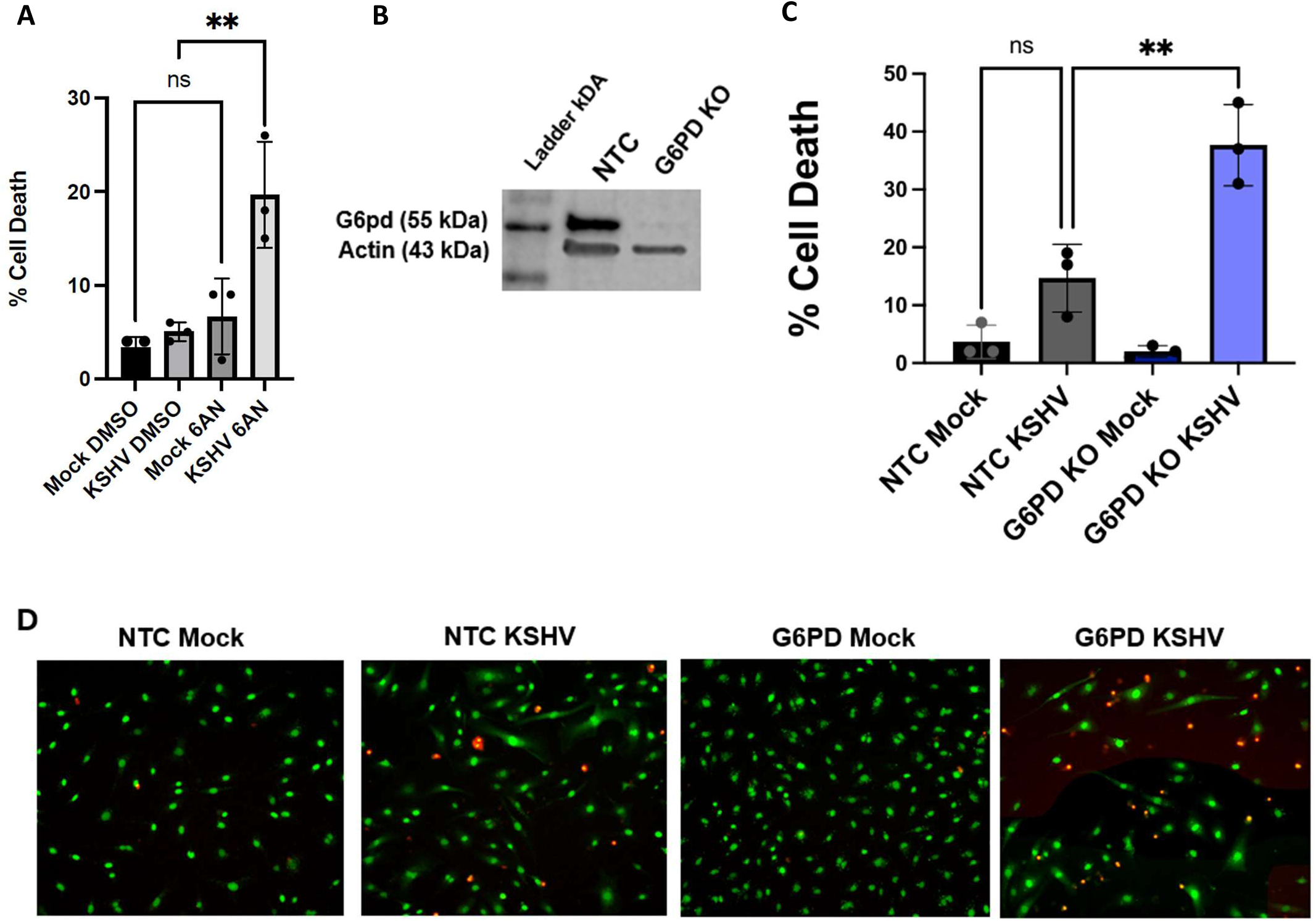
The PPP is required for KSHV Latency. (A) At 48 hours post infection, Mock and KSHV infected cells treated with either DMSO (control) or 6-AN (0.125mM), an inhibitor of the PPP enzyme glucose-6-phosphate dehydrogenase (G6PD). A trypan blue assay was performed 48 hours post drug treatment to measure cell death in all conditions. P value = 0.0045 (B). Western blot analysis of G6PD expression in both NTC and G6PD KO TIME cells. 10ug of protein was loaded for both samples and anti-actin antibody was used as a loading control. (C) TIME cells transduced with either NTC or G6PD guide RNAs and Cas9 were mock and KSHV infected. Total cell death of NTC and G6PD KO TIME cells were measured 48 hours post mock or KSHV infection. Total cell death was determined by quantifying dead cells (propidium iodide) / total cell count (SytoGreen24). P value = 0.0017 (D) Representative Images of both Mock or KSHV infection of NTC or G6PD KO TIME cells in the CellCyte live cell imager. SytoGreen24 is total cell dye which stains all cells green. Propidium iodide stains dead cells red. Data are presented as mean ± standard deviation. One way ANOVA and Tukey’s multiple comparisons test were performed for statistical significance. N =3 for all experiments.

To confirm that the cell death we see in KSHV infected cells is due to the inhibition of G6PD, and not a result of off-target effects of 6-AN, we created a genetic knockout of G6PD using a CRISPR/Cas9 lentiviral system. TIME cells were transduced with a lentivirus expressing Cas9 and either a non-targeting control (NTC) sgRNA or a G6PD targeting sgRNA. After transduction, TIME cells containing the plasmid were selected for with puromycin. G6PD was efficiently knocked out as confirmed by western blot analysis (Figure 2B). NTC and G6PD KO TIME cells were mock or KSHV infected and subsequently overlaid with complete media that contained fluorescent dyes staining all cells, SytoGreen24, or only dead cells, propidium iodide, and placed in live cell imager for 48 hours. 48 hours post infection, the knockout of G6PD had no effect on mock infected TIME cells as indicated by no change in the ratio of PI staining cells to SytoGreen staining cells (dead cells/total cells). However, there was a significant increase cell death in G6PD KO TIME cells latently infected with KSHV. On average 38% percent of the G6PD KO TIME cells stained for PI (dead cells) in comparison to NTC KHSV infected cells’ 15% of the NTC cells (Figure 2C). This result correlates with our 6-AN data (Figure 2A). Figure 2D shows representative images of the Sytogreen total cell dye (green) and Propidium iodide dead cell dye (red) for figure 2C from our Live cell imager at 48 hours post treatment. In summary, these experiments show that the PPP is necessary for the survival of endothelial cells latently infected with KSHV.

### The PPP is not required in KSHV latently infected cells to return glycolytic metabolites from the non-oxPPP to glycolysis

As shown in Figure 1A, there are two parts of the PPP, an oxidative (oxPPP) branch and a non-oxidative (non-oxPPP) branch. Glucose-6-phosphate (G6P) is the first intermediate of glucose metabolism and the connection point to the oxidative part of the PPP (42). Within this portion of the PPP, G6P is converted into ribulose-5-phosphate with the production of two NADPH molecules through a series of irreversible enzymatic reactions. The non-oxPPP branch includes multiple reversible carbon-carbon rearrangements. Ribulose-5-phosphate is the metabolite that links the non-oxidative and oxidative portions of the PPP. Ribulose-5-phosphate undergoes reversible conversions into either Xylulose 5-phosphate or ribose-5-phosphate (42)(Figure 1A). Importantly, a key enzyme, transketolase (TKT), converts non-oxPPP intermediates xylulose 5-phosphate and erythrose 4-phosphate into glycolytic intermediates fructose-6-phosphate and glyceraldehyde 3-phosphate respectively (42). The reversible nature of the non-oxPPP branch allows for the PPP to replenish either glycolysis or the PPP by creating both glycolytic and PPP intermediates depending on the cell’s metabolic needs (31).

To determine if the PPP is required in KSHV latently infected cells to increase access to glycolytic metabolites through the non-oxPPP, we tested if Oxythiamine (OT), an inhibitor of transketolase, increases cell death in latently infected (43). Mock and KSHV latently infected cells were treated with media containing either with vehicle, 5mM Oxythiamine or 10mM Oxythiamine for an additional 48 hours post establishment of latency (total of 96 hours post infection) in TIME cells. After treatment with Oxythiamine, cell death was quantified via a trypan blue exclusion assay. Neither Mock nor KSHV-infected TIME cells were sensitive to Oxythiamine (Figure 3A).

**Figure 3:**
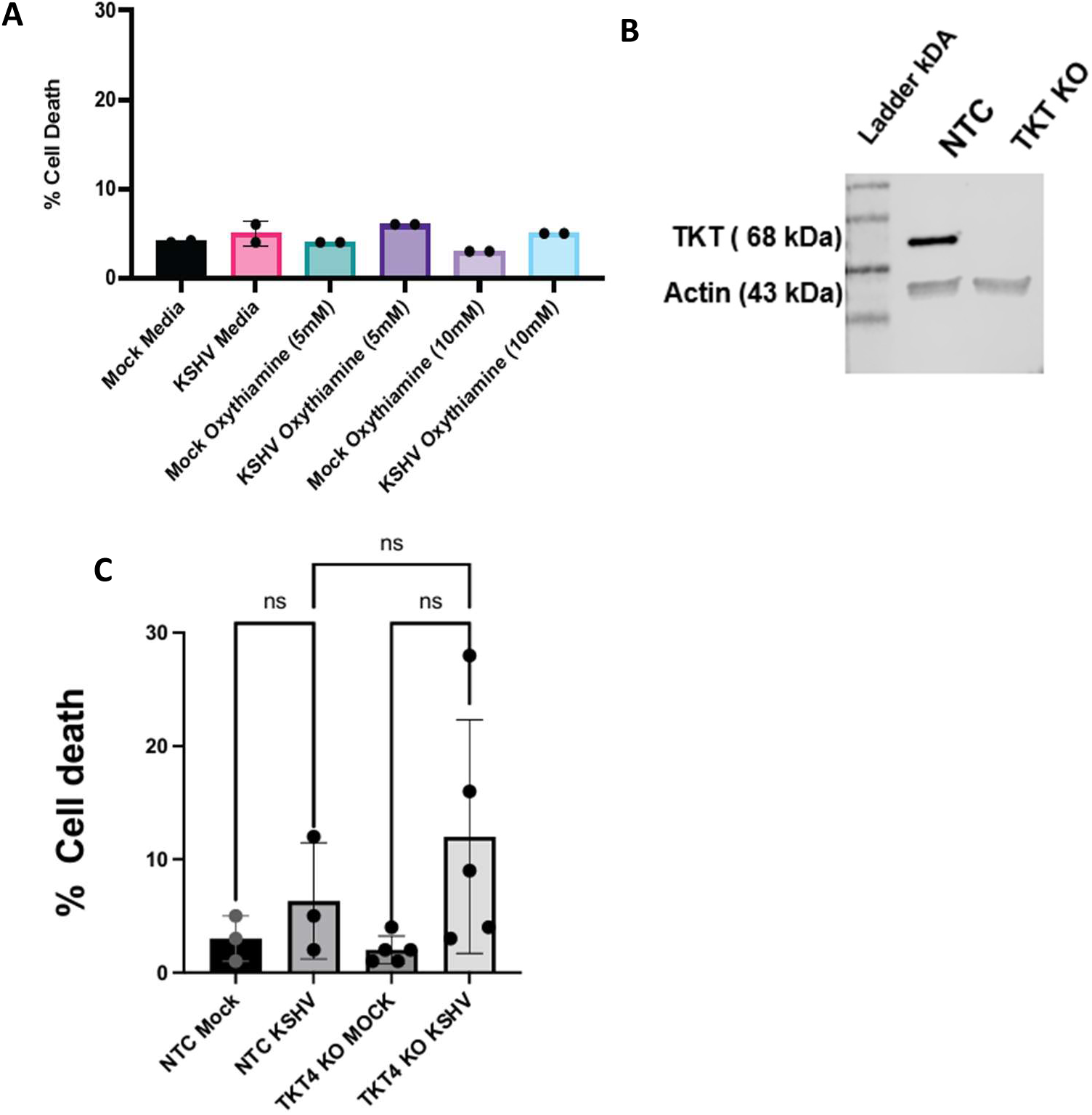
The non-oxidative PPP enzyme, Transketolase, is not necessary for the survival of KSHV latently infected cells. A) At 48 hours post infection, Mock and KSHV infected cells were treated with either media (control) or different concentrations of Oxythiamine (OT), an inhibitor of the non-oxidative PPP enzyme transketolase (TKT). A trypan blue assay was performed to measure cell death 48 hours post drug treatment. Oxythiamine does not increase cell death in Latently infected TIME cells. N=2 (B). Western blot analysis of Transketolase (TKT) expression in both NTC and TKT KO TIME cells. 10ug pf protein loaded for both samples. (C) Percent cell death in mock and KSHV infected NTC and TKO KO TIME cells 48 hours post infections determined by the ratio of dead cells (PI staining cells) to total cells (Sytogreen24). The difference in percent cell death was not statistically significant. N=3 for NTC replicates and N = 5 for TKT KO replicates. One way ANOVA and Tukey’s multiple comparisons test were performed for statistical significance.

To confirm the Oxythiamine data, we created genetic knockouts of TKT using a CRISPR/Cas9 lentiviral system as described in the previous section. TKT was efficiently knocked out as confirmed by western blot analysis (Figure 3B). NTC and TKT KO TIME cells were infected with KSHV and placed in a live cell imager for 48 hours. At 48 hours post infection, KSHV infection did not lead to a significant increase cell death in TKT KO or NTC TIME cells (Figure 3C) matching our Oxythiamine result from figure 3A. In summary, these experiments show that the cell death we see in figure 2 is not due to the inhibition of the PPP’s ability to produce glycolytic metabolites, fructose-6-phosphate and glyceraldehyde 3-phosphate.

### Purine and Pyrimidine synthesis is required for the survival of latently infected endothelial cells

There are two main byproducts of the PPP that have been shown to be advantageous for both viral infection and cancer cell proliferation, the production of 2 NADPH molecules and the production of ribose-5-phosphate, a precursor for *de-novo* nucleotide synthesis (32, 44, 45). Our previous metabolomics analysis found that both uridine and guanine were at elevated levels in KSHV-infected cells in comparison to mock infected cells (24). Intracellular levels of guanine are significantly elevated by 4.7-fold at 96 hours post infection and uridine is elevated by 2.2-fold at 96 hours post infection (Figure 4A). Uridine is a pyrimidine ribonucleoside, while guanine is a purine nucleobase.

**Figure 4:**
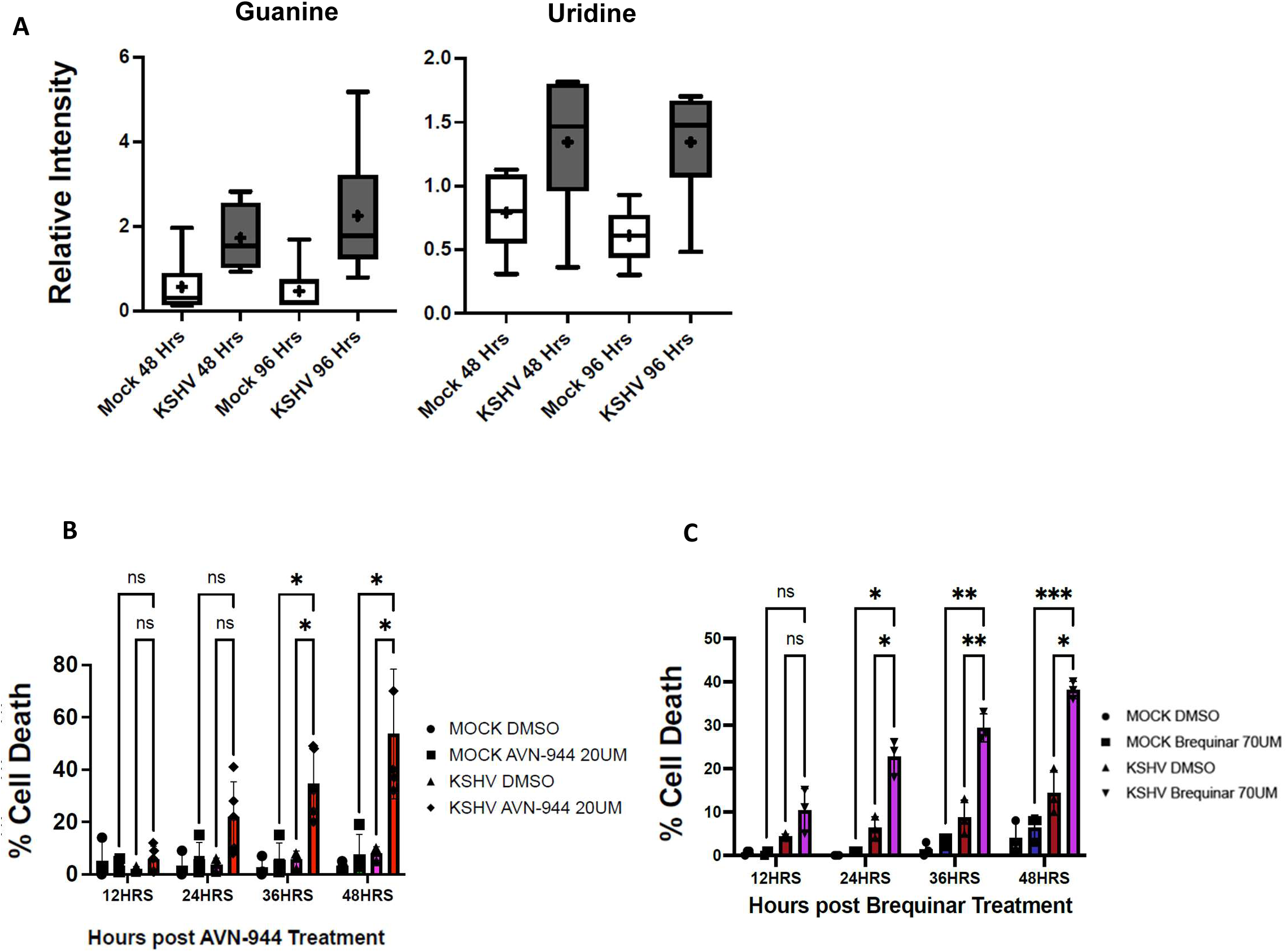
Purine and Pyrimidine nucleotide synthesis is necessary for the survival of KSHV latently infected cells. (A) Box and whisper plots of average intensity of nitrogenous purine base guanine and pyrimidine nucleoside uridine as determined by global metabolomics screen of Mock and KSHV infected TIME cells at 48 and 96 hours post infection. Mock infected cells are represented by white boxes while KSHV infected cells are represented in grey. Measurements were taken from 6 biological replicates. The average of each group is represented by a (+) sign. P values ≤ 0.05. (b) 48 hours post infection, Mock and KSHV infected cells were treated with either DMSO (control) or AVN-944, an inhibitor of Inosine-5′-monophosphate dehydrogenase (IMPDH), for an additional 48 hours. Total cell death was measured by propidium iodide and total cell count was measured by SytoGreen24 over the course of the drug treatment in a live cell imager. Percent cell death = dead cells (PI staining cells) divided by total cells (Sytogreen24). Two way ANOVA and Tukey multiple comparisons test, n=3, P value 36 Hours* = 0.02, P value 48 Hours* = 0.04 (C) 48 hours post infection, Mock and KSHV infected cells were treated with either DMSO (control) or dihydroorotate dehydrogenase (DHODH) inhibitor, Brequinar. Percent cell death measured over the course of the 48-hour drug treatment as stated in b. Two-way ANOVA and Tukey multiple comparisons test, n=3, P value 24 Hours * ≤ 0.03, P value 36 Hours ** ≤ 0.009, P value 48 Hours* = 0.02, **=0.001, *** = 0.0007.

To distinguish if de novo synthesis of purines are necessary for KSHV latency, we assessed the effect of AVN-944. AVN-944 is an inosine 5’-monophosphate dehydrogenase (IMPDH) inhibitor (46). IMPDH is upstream of guanine monophosphate (GMP). Inhibition of IMPDH leads to a decrease in intracellular guanine monophosphate (46). At 48 hours post infection, mock and KSHV infected cells were treated with DMSO or AVN-944 (20uM) for an additional 48 hours. Cell death was tracked in a live cell imager over the course of the drug treatment. Starting from 24 hours post drug treatment, KSHV-infected TIME cells were more sensitive to AVN-944 treatment in comparison to Mock-infected cells (Figure 4B). After 48 hours post drug treatment, there was a large time dependent increase in cell death in KSHV-infected cells where on average 54% of KSHV infected cells died due to treatment of AVN-944 in comparison to mock’s 8% cell death (Figure 4B).

Brequinar is an inhibitor of the pyrimidine synthesis enzyme dihydroorotate dehydrogenase (DHODH). Inhibition of DHODH blocks the formation of orotate, which is essential for *de-novo* synthesis of pyrimidines and leads to a decrease in the intracellular pool of pyrimidines (47). Mock and KSHV infected TIME cells were treated with brequinar similarly to AVN-944 as just described. At 24 hours post drug treatment, KSHV-infected TIME cells were more sensitive to brequinar treatment in comparison to Mock cells (Figure 4C). By 48 hours post drug treatment, basal cell death levels had rose in KSHV-infected TIME cells while their mock counterparts remained low. Vehicle treated KSHV infected TIME cells showed 13% cell death while the cell death in the KSHV Infected TIME cells treated with brequinar approached 40% (Figure 4C). Taken together, these experiments show that both purine and pyrimidine synthesis is necessary for the survival of endothelial cells latently infected with KSHV.

### The PPP is required for nucleotide synthesis in KSHV latently infected TIME cells

To determine if the cell death we see with the inhibition of the PPP with 6AN in Figure 2 is related to a reduction in nucleotides, we tested if KSHV latently infected cells could be rescued with the addition of D-ribose. Exogenous D-ribose can be converted to ribose-5-phosphate by the enzyme ribokinase, therefore, D-ribose can provide a source of ribose-5-phosphate (48).

Mock and KSHV infected TIME cells were treated with 6-AN and two different concentrations of D-ribose. As seen in figure 2, KSHV-infected TIME cells were more sensitive to 6-AN showing an average 23% percent cell death in comparison to Mock cells’ 2% (Figure 5A). Addition of D-ribose at 3.5mM or 7mM had no significant effect on Mock or KSHV infected cells treated with DMSO. There is a dose dependent response of KSHV-infected cells treated with 6AN and D-ribose. There is a trend for rescue of 6-AN induced cell death in KSHV infected cells with D-ribose at 3.5mM. However, the addition of D-ribose at 7.0mM leads to a significant, albeit incomplete, rescue of cell death in KSHV-infected cells treated with 6AN (Figure 5A). These results indicate that at least part of the necessity of the PPP in KSHV latently infected TIME cells comes from the requirement for ribose-5-phosphate and therefore, nucleotide synthesis as the next steps in the non-oxPPP (Figure 1A) are not necessary for KSHV latency.

**Figure 5:**
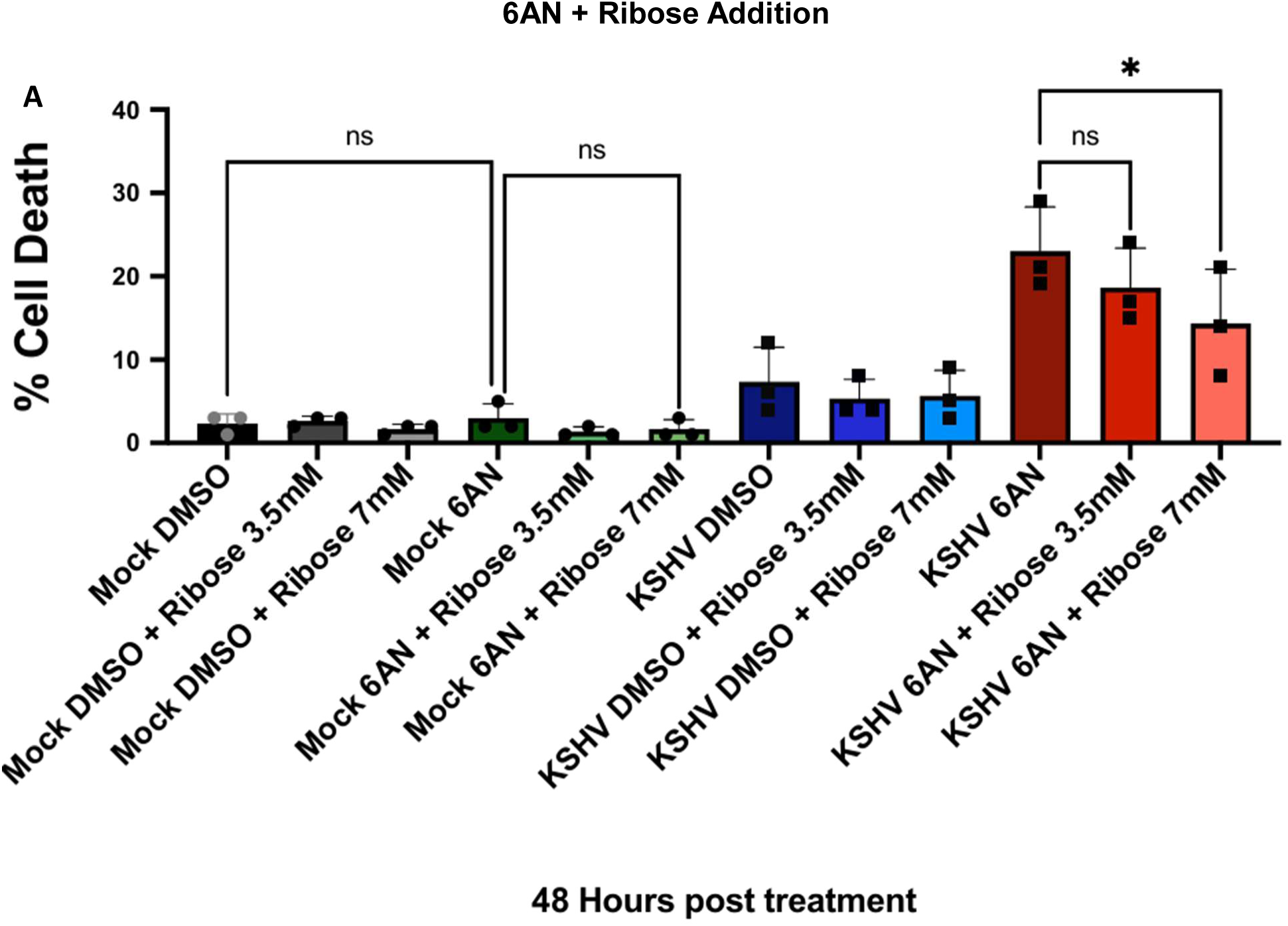
The PPP is required for nucleotide synthesis in KSHV latently infected cells. (A) TIME cells were mock or KSHV-infected. At 48 hours post infection, Mock and KSHV infected cells treated with either DMSO (control), 6-AN (0.3mM) or 6-AN (0.3mM) and D-ribose at 3.5mM or 7mM. In addition to the drug treatment, all conditions received SytoGreen24 (Total cell dye) and Propidium iodide (dead cell dye). Cell death was measured over the course of the drug treatment in a live cell imager. Cell death at 48 hours post drug treatment was measured and represented as a bar graph. Data are presented as means. One way ANOVA and sidaks’s multiple comparisons test were performed for statistical significance. N =3 for all experiments. P value * = 0.02.

## Discussion

In normal endothelial cells, a low percentage of glucose, approximately 1-3% (49, 50), is shunted to the PPP. In many cancer cells, as well as cells infected with lytic viruses, the PPP flux increases and in endothelial cells the PPP can be increased during angiogenesis (33, 51–53). Here we show that the oncogenic virus, KSHV, increases metabolites in the oxidative half of the PPP during latent infection of endothelial cells. We have previously shown that KSHV induces glucose uptake and glycolysis which could support the increased PPP (24, 25).

Interestingly, latent KSHV infection of TIME cells is dependent on the PPP for survival as chemical inhibition or knockout of the first and rate-limiting step of the PPP, G6PD, leads to significantly increased cell death in latently infected TIME cells but not mock infected cells. Therefore, the increase in PPP intermediates during infection is likely increased carbon flow through the PPP and not a block downstream of the upregulated metabolites.

The PPP is made up of two halves, the oxidative half and the non-oxidative half. The oxidative half is not reversible and produces NADPH and ends in the production of ribulose-5-phosphate. The non-oxidative half of the PPP produces fructose-6-phosphate (F-6-P) and glyceraldehyde-3-phosphate (G3P) in reversible steps that can be shunted back into glycolysis. Inhibiting G6PD knocks out the oxidative half of the PPP. TKT is an enzyme important for two steps of the non-oxidative half of the PPP. Chemical inhibition or knockout of this enzyme has no effect on latently infected cells demonstrating that the oxidative half of the PPP is required for survival of latently infected cells but not the non-oxidative half. Therefore, KSHV latent infection is not dependent on flux back into glycolysis from the non-oxidative half of the PPP.

The addition of ribose to latently infected cells can recover much of the cell death from 6-AN treatment. As ribose can be converted into ribose-5-phosphate, it appears that ribose-5-phosphate is a novel KSHV dependency factor made by the PPP. Because the non-oxidative half of the PPP is not required, conversion of ribose-5-phosphate to G3P or F6-P for glycolysis is not the essential role of the PPP during KSHV latency. Therefore, it is highly likely that the PPP and ribose-5-phosphate is required for nucleotide synthesis. In alignment with this hypothesis, we showed that both de-novo purine and pyrimidine synthesis are required for the survival of KSHV latently infected TIME cells as inhibition of either pathway led to significant increased cell death in latently infected cells as compared to mock infected cells. Taken together, the PPP appears to be required for nucleotide synthesis during latent KSHV infection.

Many lytic viruses have been shown to be dependent on the PPP and nucleotide synthesis including gamma herpesviruses. MHV-68 and CMV were shown to require the PPP and nucleotide synthesis for lytic replication (45, 54). Epstein–Barr virus (EBV), another oncogenic ɣ-herpesvirus, has been shown to alter nucleotide synthesis during lytic replication and inhibition of pyrimidine synthesis in EBV leads to impaired lytic replication (55). EBV induces the production of nucleotide cytidine-5’-triphosphate (CTP) and other segments of the de-novo pyrimidine synthesis pathway through NFκB, EBNA2, and MYC (55). This upregulation is also necessary for the survival of EBV transformed B-cells (55). KSHV was also previously shown to require nucleotide synthesis for lytic replication (56). There is also recent evidence that supports that nucleotide synthesis is necessary for KSHV latency. The Latent viral gene vCyc interacts with CAD an enzyme within the de novo pyrimidine synthesis pathway to promote pyrimidine synthesis during lytic replication (51). This activated version of CAD can also promote aerobic glycolysis through deamination of NF-κB (51). Further work is necessary to determine if vCyc or other latent genes induce the requirement for the PPP during latent infection.

While these examples show how nucleotide synthesis can be necessary for lytic replication, the question of why nucleotide synthesis would be necessary during latent infection remains. Since KSHV is not rapidly replicating its genome during latency and does not induce rapid cell proliferation of cultured endothelial cells, nucleotide synthesis may be required for other metabolic pathways or for specific cell signaling pathways during latent infection. Angiogenesis also alters the metabolic state of endothelial cells (57). We have previously shown that KSHV induces angiogenic phenotypes during latent infection, including increased motility, which may require different metabolic pathways including the PPP and nucleotide synthesis (33, 58). That the PPP and nucleotide synthesis is required during latent infection raises the possibility that lytic viruses do not require the induction of the PPP solely for genome replication. The PPP and nucleotide synthesis may also be required to keep the cells alive to allow replication to be completed.

While the PPP appears to be necessary for nucleotide synthesis during latent KSHV infection of endothelial cells, the addition of ribose did not completely protect latently infected cells treated with the PPP inhibitor, 6-AN. While the amount of ribose that enters the cell is unknown, it may not be enough to recover the phenotype. However, it is also possible that other factors from the PPP are required. The oxidative half of the PPP produces NADPH which is required for fatty acid synthesis (31). We previously showed that KSHV induced most of the long chain fatty acids in latently infected cells and that inhibitors of fatty acid synthesis led to increased cell death in latently infected endothelial cells (24). NADPH is also needed to resolve reactive oxygen species by interacting with cellular antioxidant enzymes, glutathione (GSH) and thioredoxin (59). KSHV latent infection has also been shown to increase reactive oxygen species (60). Therefore, both nucleotide synthesis and NADPH could be additional dependency factors produced by the PPP during latent infection.

Viruses have developed many strategies to obtain the necessary machinery from their host cell for infected cell survival. One of these strategies is altering host cell metabolic pathways to meet the specific needs of the virus to ensure a successful infection. Like cancer cells, oncogenic viruses need to alter metabolic processes to proliferate and survive. Inhibitors of the PPP could be candidates for novel treatments of latent KSHV infection of endothelial cells and ultimately KS tumors. There have been multiple efforts to target nucleotide synthesis by inhibiting different critical enzymes within nucleotide metabolism or creating nucleotide analogues to mimic natural nucleotides in order to block DNA or RNA synthesis within infected cells or cancer cells (61, 62). These inhibitors could be potential therapeutic avenues for KSHV and KS. By targeting cellular dependency factors in latently infected cells, the common issue of relapse after standard chemotherapy might be alleviated. Further work will be needed to determine the efficacy of these inhibitors in vivo.

## Materials and Methods

### Cell lines

Tert-immortalized microvascular endothelial (TIME) cells (34) were maintained as a monolayer in endothelial cell growth medium-2 (EGM-2) media (Lonza) supplemented with an EGM-2 MV BulletKit (Lonza) containing 5% fetal bovine serum (FBS), hydrocortisone, ascorbic acid, Insulin-like growth factor, vascular endothelial growth factor, ascorbic acid, and human endothelial and fibroblast endothelial growth factors. 293T cells were maintained in Dulbecco’s modified Eagle media (DMEM) supplemented with 4.5g/L glucose, 10% FBS, 1% L-glutamine, 1% sodium pyruvate and 1% penicillin-streptomycin antibiotic mix.

For virus production, BCBL-1 cells (35) were grown in Roswell Park Memorial Institute media (RPMI) 1640 supplemented with 10% FBS, L-glutamine, penicillin-streptomycin, and 2-mercaptoethanol. For BAC16 virus production, iSLK cells were grown in DMEM supplemented with 4.5g/L glucose, 10% FBS, L-glutamine, sodium pyruvate, penicillin-streptomycin, 250μg/mL G418, 1200μg/mL hygromycin B, and 1μg/mL puromycin (36).

### Reagents and antibodies

6-Aminonicotinamide (6-AN) (Sigma) was diluted in DMSO and used at final concentrations 0.125mM and 0.3mM. Oxythiamine (Sigma) was diluted in water and was used at final concentrations 5mM and 10mM. AVN-944 (Medchem) was diluted in DMSO and used at final concentration of 20uM. Brequinar (medchem) was diluted in DMSO and used at final concentration of 75uM. D-Ribose (Sigma) was diluted in water and used at final concentrations of 3.5mM and 7mM. Green-fluorescent nucleic acid stain SYTO 24 Green (Invitrogen) was diluted in DMSO and used at final concentration of 2uM. Propidium iodide (PI) was diluted in water and used at a final concentration of 2ug/mL.

Antibodies G6PD (ABclonal #A13983), and TKT (Abclonal #A27144) were used at 1:1000 dilution while β-actin (Sigma # A5441) was used at 1:10,000 as specified by the manufacturer. Primary antibodies that recognize the KSHV protein, LANA, was used at 1:1000 dilution.

### Virus Infections and Immunofluorescence

5 × 10^5^ cells/mL BCBL-1 cells were induced with 20ng/mL of TPA (12-O-tetradecanoylphorbol-13-acetate; Sigma) to produce KSHV inoculum as described previously (16). 5-6 days post induction, cells were pelleted, and the supernatant was run through a 0.45μm-pore-size filter (Whatman) to remove cell debris. Virions were pelleted at 30,000xg for 2 hours at 4°C in a JA-14 rotor, Avanti-J-25 centrifuge (Beckman Coulter). The viral pellet was resuspended in EBM-2 without supplements. The KSHV Bac16 viruses were made as described previously (37, 38).

KSHV infection of all cell types was performed in serum-free endothelial cell growth media (Lonza or PromoCell) supplemented with 8ug/mL polybrene for 4 hours. Post infection, the media was replaced with complete EGM-2 media (Lonza) after washing cells with phosphate-buffered saline (PBS). Mock infections were performed identical to KSHV infections without the addition of concentrated virus to the serum-free media. To measure infection rates, an aliquot of mock or KSHV infected TIME cells was seeded on Corning Falcon 4-well tissue culture-treated slides (VWR) and fixed with 4% (wt/vol) paraformaldehyde in PBS. Briefly, fixed cells were blocked using an immunofluorescence blocking buffer (1% BSA in TBS) then incubated with a primary antibody for a latent KSHV protein, LANA, for 1 hour. Cells were then incubated with a fluor-conjugated secondary antibody (Molecular Probes/Invitrogen) at a dilution of 1:1,000 for 1 hour. Before being viewed under a Keyence BZ-X710, Cells were mounted in medium containing DAPI (4’,6’-diamidino-2-phenylindole). Infections where at least 88% of cells expressed LANA were used for experiments.

### CRISPR/Cas9 gene targeting, Transfection and Transduction

To generate lentiviruses to knock out G6PD or TKT, guide RNAs (gRNAs) specific to each of these genes were obtained from a human CRISPR Metabolic Gene Knockout Library (39). Sense and antisense single gRNAs specific to each gene were annealed to form a gRNA. Each gRNA was annealed into a pRRL plasmid expressing a Cas9-T2A cassette (a gift from Daniel Stetson, University of Washington) using an in-Fusion cloning system (Takara Bio). Cloned plasmids were then transformed into Stellar Competent E. coli cells and selected for on LB plates with carbenicillin. LB broth with carbenicillin was inoculated with a single colony and incubated overnight at 37 °C in a shaking incubator. Plasmids were isolated using a Qiagen Mini-Prep Kit. Non-targeting control (NTC) gRNAs plasmids were created identical to the knockout gene gRNAs, however NTC gRNAs do not have any human gene target.

List of sgGRNAs:

sgG6PD_2S: GCAAGGCCAGGTAGAAGAGG

sgG6PD_2AS: CCTCTTCTACCTGGCCTTGC

sgTKT_4(S): GGGCACCGTCCTGTTGCGGG

sgTKT_4(AS): CCCGCAACAGGACGGTGCCC

NTC_A_267(F): GTTCGATCCGGCGACTCGAA

NTC_A_267(R): TTCGAGTCGCCGGATCGAAC

NTC_B_291(F): GACATTGTTAGTAACGACTC

NTC_B_291(R): GAGTCGTTACTAACAATGTC

Lentiviruses targeting G6PD, TKT or the non-targeting control was generated by co-transfection of 293T cells. 5.0 x 10^6^ HEK293T cells were seeded per a 10 cm dish the night before transfection. Transit 293T (Mirus Bio) was used to transfect plasmids (8μg of psPAX2, 4μg of PMD2.G, and 8μg of the pRRL vector) as indicated by the manufacturer. 24 hours post transfection, cells were given fresh complete DMEM Media. Media containing lentivirus was collected at 48 and 72 h post transfection and filtered through 0.45-μm filters before aliquoting and freezing at −80°C.

Lentiviral infections were done by incubating cells with 1mL of virus for 6-well plates or 1.5mLs for T25 flask for 6 hours in the presence of 8μg/mL polybrene. Transduced cells were treated with puromycin (1 μg/mL) selection for 2-3 days. After selection, cells were allowed to grow out for 3-10 days in fresh complete media.

### Western blot analysis

Adherent cells were trypsinized, harvested and pelleted before being washed with PBS. Cell pellets were lysed with RPIA lysis buffer (50mM Tris-HCl, pH 7.6, 150 mM NaCl, 1 mM EDTA, 1% Nonidet P-40, 0.5% deoxycholate, 0.1% sodium dodecyl sulfate, 1 mM sodium orthovanadate, 1 mM sodium fluoride, 40 mM β-glycerophosphate, Roche Complete Mini protease inhibitor tablet). Protein concentrations were determined by using a Pierce BCA assay (Thermo Fisher Scientific) on cell lysates. Equal amounts of protein were loaded into a 4–20% polyacrylamide gel (BioRad) and then transferred to a Immun-Blot PVDF Membrane (BioRad) after separation on the polyacrylamide gel. Membranes were blocked with TBS or PBS Blocking Buffer (LICORbio) for 1 hour at room temp and then blotted with primary antibodies diluted in TBS buffer overnight at 4°C. Dilutions for antibodies were stated above. Blotted membranes were washed three times for 10 mins at room temp with TBS-T then treated with LI-COR IRdye secondary antibodies, IRDye 680RD and IRDye 800CW, for 1 hour at room temperature. After treatment, membranes were washed with TBS-T and imaged on the Odyssey Fc system.

### Inhibitor and Ribose Addition Assay

For drug inhibition assays, 2.0 x 10^5^ - 2.2 x 10^5^ TIME cells were seeded in 6-well plates and were mock or KSHV infected. 48 hours post infection, cells were treated with complete media that contained the drug inhibitor, SYTO 24 green (green fluorescent total cell dye) and Propidium iodide (PI) (red fluorescent cell death dye) at the concentrations stated above for an additional 48 hours post infection. These plates were placed into a CellCyte live cell imager (Cytena) where fluorescent images were taken every 12 hours for the entire course of the drug treatment. Images taken for the duration of the assay were used to assess the percentage of dead cells by using the CellCyte analysis software to count total cells and total dead cells per treatment.

Ribose addition experiments were performed similar to drug inhibition assays however, cells were treated with complete media that contained D-ribose, 6-AN, SYTO 24 green and Propidium iodide (PI) for an additional 48 hours post infection. These plates were also placed into a CellCyte live cell imager (Cytena) where fluorescent images were taken every 12 hours and percentage of dead cells was measured using the CellCyte analysis software to count total cells and total dead cells per treatment.

### Trypan Blue Assay

For some experiments, cell death was measured via a trypan blue assay. Before TIME cells were trypsinized, complete media containing dead cells and PBS used to wash the cells were collected together. Trypsinized cells were then added to that pool of cells before the entire pool was pelleted down. Pelleted cells were resuspended in 50-100uL of complete media. 15uL of resuspend cells were added to equal amounts trypan blue, loaded into a chamber slide and counted by a TC-20 cell Counter (Bio-rad).

### Statistical analysis

Data and Statistical analysis was done using GraphPad PRISM software.

## Data availability

All data from this study is included in this article itself.

